# Discovery and Optimization of Imidazothiazole Carboxamides as Novel Anti-Tuberculosis Agents

**DOI:** 10.64898/2026.09.21.753226

**Authors:** Leo Meng, David Grant, Han Xie, Kirsten Tolentino, Rachel Ford, Jasmine Webb, Case W. McNamara, Baiyuan Yang, Arnab K. Chatterjee

**Author notes:** Corresponding author: Baiyuan Yang. These authors contributed equally to this work: Leo Meng, David Grant, Han Xie.

## Abstract

Screening of the open-access CRESTdb small-molecule library identified the imidazothiazole carboxamide **sCQG20**0 (**1**) as an initial hit, with activity against H37Rv and Erdman *Mycobacterium tuberculosis* strains in cholesterol-containing medium (MIC_90_= 6.88 µM and 4.48 µM, respectively) and against intracellular Mtb (EC_50_= 3.80 µM). **sCQG200** retained activity against drug-resistant Mtb strains, including clinical MDR/XDR isolates. SAR optimization led to analogs **21** and **24** with substantially improved antimycobacterial potency. Mouse pharmacokinetic studies following oral and intravenous administration showed 62.1% and 29.5% oral bioavailability for **21** and **24**, respectively. This structurally differentiated chemotype provides a promising starting point for further optimization as antitubercular agent.

---

Tuberculosis (TB), caused by *Mycobacterium tuberculosis* (Mtb), remains the world’s leading cause of death from a single infectious agent.^1^ Drug-susceptible TB is generally treated with multidrug therapy for 4–6 months, with a 4-month regimen now recommended for eligible patients with rifampicin-susceptible TB, while the conventional 6-month regimen remains widely used.^1,2^ Multidrug-resistant tuberculosis (MDR-TB), defined by resistance to at least isoniazid and rifampicin, and rifampicin-resistant tuberculosis (RR-TB) require alternative multidrug regimens whose duration and composition depend on the resistance profile and eligibility for shorter regimens.^2,3^ MDR/RR-TB remains particularly challenging because drug resistance limits the effectiveness of standard regimens and may necessitate more complex treatment. Despite recent advances in shorter, all-oral regimens, treatment of drug-resistant TB remains associated with substantial treatment burden and potential drug-related toxicity.^2,3^ Drug-resistant TB is estimated to account for approximately 13% of antimicrobial-resistance-attributable deaths worldwide.^4^ Collectively, these challenges highlight the continued need for new anti-TB agents with novel mechanisms of action (MoAs) capable of shortening treatment duration, improving tolerability, and retaining activity against drug-resistant strains.

Phenotypic high-throughput screening under growth conditions incorporating host-relevant carbon sources provides a complementary approach for identifying novel antitubercular chemical matter. During infection, Mtb can utilize host-derived cholesterol as a source of carbon and energy, and cholesterol uptake and catabolism contribute to bacterial survival and persistence in the host.^5,6^ Accordingly, screening compounds in cholesterol-containing medium can reveal conditional vulnerabilities that may not be readily detected under conventional nutrient-rich growth conditions. Previous chemical screens at Calibr have identified structurally diverse compounds that inhibit Mtb growth in cholesterol-containing medium.^7-11^ Such approaches can therefore provide access to chemical matter with distinct mechanisms of action and expand the chemical and mechanistic diversity of the TB drug discovery pipeline.

As part of our continued efforts to identify new chemical entities for TB treatment, we conducted a primary phenotypic screen against *Mycobacterium tuberculosis* (Mtb) H37Rv in cholesterol-supplemented 7H12 medium using a 384-well format. Approximately 24,000 novel compounds from our open-access CRESTdb library^12^ were screened at a concentration of 10 μM. The library contains approximately 240 distinct chemical series, with up to ∼100 analogs represented within selected series, enabling rapid exploration of the surrounding chemical space and efficient structure–activity relationship assessment of confirmed screening hits. Additionally, common building blocks are also available, mostly on a multigram scale, facilitating rapid medicinal chemistry optimization once a hit is identified. Through this screening, we identified **sCQG200 (1)** as a top hit, exhibiting activity against *M. tuberculosis* grown in cholesterol-containing medium, with an H37Rv MIC_90_ of 6.88 μM. **sCQG200** also showed activity against Erdman strain and intracellular Mtb, with an MIC_90_ and EC_50_ of 4.48, 3.8 μM respectively. The observed anti-Mtb activity was unlikely to result from nonspecific cytotoxicity, as **sCQG200** exhibited CC_50_ values >40 μM in both mammalian cell lines HEK293T and HepG2 cells (**Table 1**). More importantly, **sCQG200** retained activity against clinical extensively drug-resistant (CXDR) *M. tuberculosis* isolates, with MIC_90_ values of 3.12 µM against CXDR-1 and 12.5 µM against CXDR-2, as well as against a clinical multidrug-resistant (CMDR) isolate (MIC_90_ = 3.12 µM) (**Table 1**). The retention of activity against these drug-resistant isolates suggests that **sCQG200** is not substantially affected by the resistance mechanisms represented in this panel and supports this chemotype as a differentiated starting point for further optimization. Structurally, sCQG200 features an imidazothiazole carboxamide chemotype comprising an aryl substituent, a rigid fused N/S-containing heteroaromatic core, and a carboxamide functionality. The imidazothiazole scaffold has established precedent in medicinal chemistry, as exemplified by tetrahydroimidazothiazole-containing drug levamisole (**Figure 1**). Imidazo[2,1-b]thiazole-5-carboxamides including ND-11543 have also been reported as potent antitubercular agents targeting QcrB,^13,14^ a component of the mycobacterial cytochrome bcc-aa_3_ respiratory complex (**Figure 1**). Considering that **sCQG200** also contains an imidazothiazole core, we evaluated its activity against *M. tuberculosis* strains harboring QcrB mutations associated with resistance to Q203, a known QcrB inhibitor. Notably, **sCQG200** retained activity against three Q203-resistant mutant strains, including Q203 R1 (QcrB_*T313I*_), Q203 R2 (QcrB_*M342A*_), and Q203 R3 (QcrB_*M342T*_), with an MIC_90_ of 6.25 µM against each strain, comparable to the WT MIC measured in the same assay (**Table 1**). These results suggest that **sCQG200** is unlikely to share the same QcrB-dependent mechanism of action as Q203. Taken together, these findings support sCQG200 as a structurally differentiated imidazothiazole carboxamide chemotype and a promising starting point for further SAR optimization and mechanism-of-action studies. Herein, we report detailed structure–activity relationship (SAR) studies together with ADME and pharmacokinetic characterization of this series.

**Table 1.** Antitubercular activity and cytotoxicity of the initial hit compound **sCQG200** (**1**).

|  | **sCQG200** |
| --- | --- |
| H37Rv Cholesterol MIC<sub>90</sub> (μM) | 6.88 |
| Erdman Cholesterol MIC<sub>90</sub> (μM) | 4.48 |
| H37Rv Intramacrophage THP-1 EC<sub>50</sub> (μM) | 3.80 |
| CXDR-1 / CXDR-2 MIC<sub>90</sub> (μM) | 3.12 / 12.5 |
| Q203 R1 (QcrB<sub>T313I</sub>) MIC<sub>90</sub> (μM) | 6.25 |
| Q203 R2 (QcrB<sub>M342A</sub>) MIC<sub>90</sub> (μM) | 6.25 |
| Q203 R3 (QcrB<sub>M342T</sub>) MIC<sub>90</sub> (μM) | 6.25 |
| HEK293T / HepG2 CC<sub>50</sub> (μM) | > 80 / > 80 |

**Table 1.**
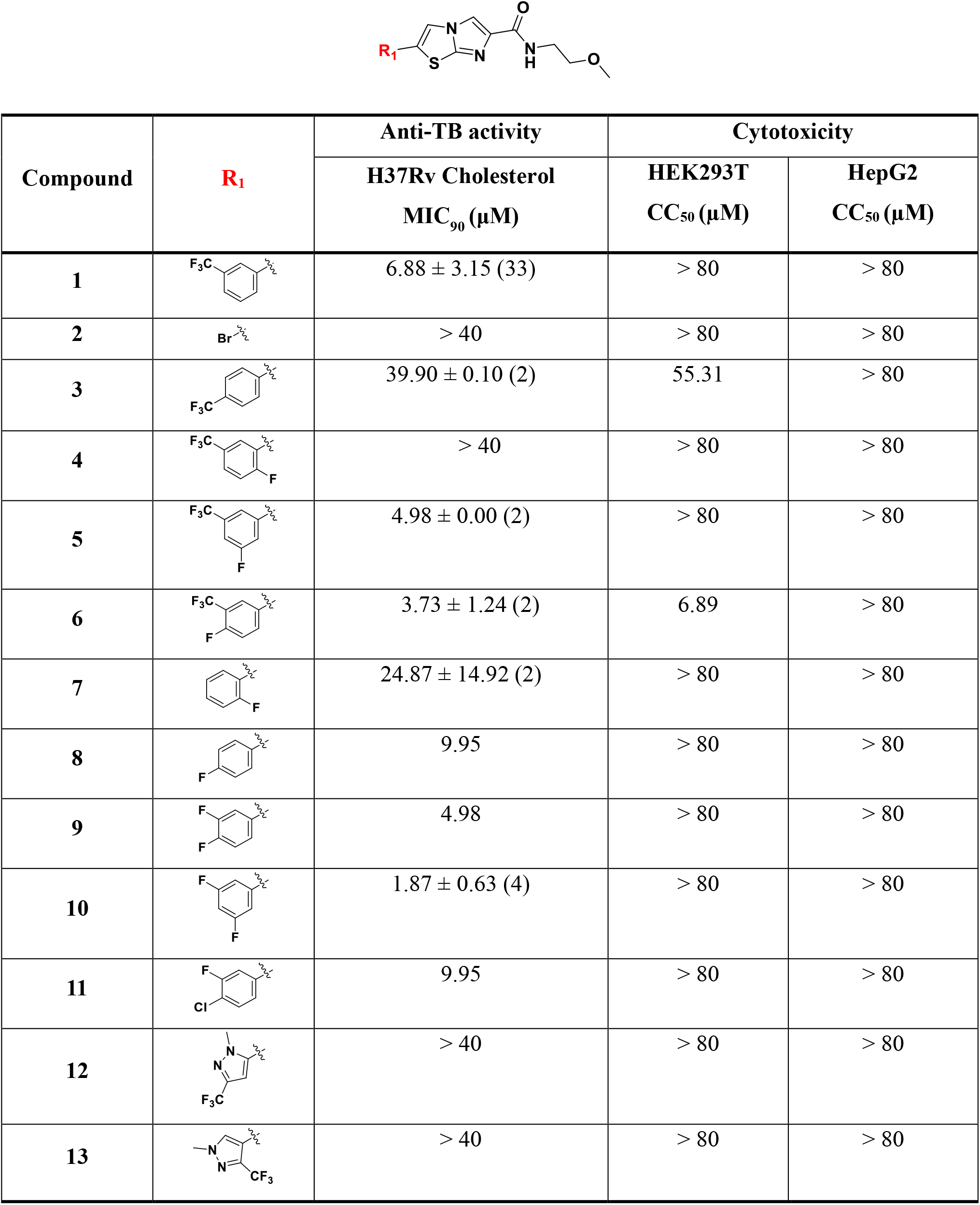
*In vitro* anti-TB inhibitory activity and cytotoxicity of aryl analogs **1 – 13**.

| Compound | **R<sub>1</sub>** | Anti-TB activity | Cytotoxicity |  |
| --- | --- | --- | --- | --- |
|  |  | H37Rv Cholesterol<br>MIC<sub>90</sub> (μM) | HEK293T<br>CC<sub>50</sub> (μM) | HepG2<br>CC<sub>50</sub> (μM) |
| **1** |  | 6.88 ± 3.15 (33) | > 80 | > 80 |
| **2** |  | > 40 | > 80 | > 80 |
| **3** |  | 39.90 ± 0.10 (2) | 55.31 | > 80 |
| **4** |  | > 40 | > 80 | > 80 |
| **5** |  | 4.98 ± 0.00 (2) | > 80 | > 80 |
| **6** |  | 3.73 ± 1.24 (2) | 6.89 | > 80 |
| **7** |  | 24.87 ± 14.92 (2) | > 80 | > 80 |
| **8** |  | 9.95 | > 80 | > 80 |
| **9** |  | 4.98 | > 80 | > 80 |
| **10** |  | 1.87 ± 0.63 (4) | > 80 | > 80 |
| **11** |  | 9.95 | > 80 | > 80 |
| **12** |  | > 40 | > 80 | > 80 |
| **13** |  | > 40 | > 80 | > 80 |

**Table 3.** *In vitro* anti-TB inhibitory activity and cytotoxicity of amide analogs **14 – 24**.

| Compound | $R_2$ | Anti-TB activity | Cytotoxicity | |
| --- | --- | --- | --- | --- |
|  |  | H37Rv Cholesterol<br>MIC<sub>90</sub> (μM) | HEK293T<br>CC<sub>50</sub> (μM) | HepG2<br>CC<sub>50</sub> (μM) |
| **10** |  | 1.87 ± 0.63 (4) | > 80 | > 80 |
| **14<sup>a</sup>** |  | > 40 | > 80 | > 80 |
| **15<sup>b</sup>** |  | 19.90 | nd | nd |
| **16** |  | > 40 | 62.28 | > 80 |
| **17** |  | 19.95 ± 0.05 (2) | > 80 | > 80 |
| **18** |  | 1.24 ± 0.00 (2) | > 80 | > 80 |
| **19** |  | 1.86 ± 0.62 (2) | > 80 | > 80 |
| **20** |  | 1.24 ± 0.002 (2) | > 80 | > 80 |
| **21** |  | 0.94 ± 0.31 (2) | > 80 | > 80 |
| **22** |  | 0.93 ± 0.31 (2) | > 80 | > 80 |
| **23** |  | 2.49 ± 0.00 (2) | > 80 | > 80 |
| **24** |  | 1.62 ± 0.64 (12) | > 80 | > 80 |

<sup>a</sup>: structure of **14**
<sup>b</sup>: structure of **15**
nd = not determined

**Figure 1.**
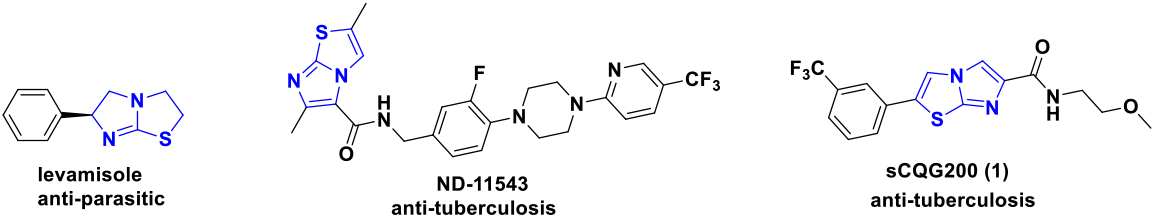
Representative imidazothiazole-derived compounds. Levamisole is an antiparasitic agent, **ND-11543** is a reported antitubercular compound as QcrB inhibitor, and **sCQG200** (**1**) is the antitubercular hit identified in this study.

A library of analogs modified at the aryl and amide regions was prepared according to the synthetic route outlined in **Scheme 1**. The synthesis commenced with hydrolysis of commercially available ethyl 2-bromoimidazo[2,1-b]thiazole-6-carboxylate (**S1**) to afford the corresponding carboxylic acid **S2**, followed by HATU-mediated amide coupling to provide amide intermediate **S3**. Subsequent Suzuki coupling furnished **sCQG200** and the related analogs. Six common building blocks for this series were retrieved from the CRESTdb intermediate stock, enabling rapid SAR optimization. All compounds were evaluated for activity against *M. tuberculosis* H37Rv using an MIC_90_assay, together with cytotoxicity assessment in HEK293T and HepG2 mammalian cell lines.

**Scheme 1.**
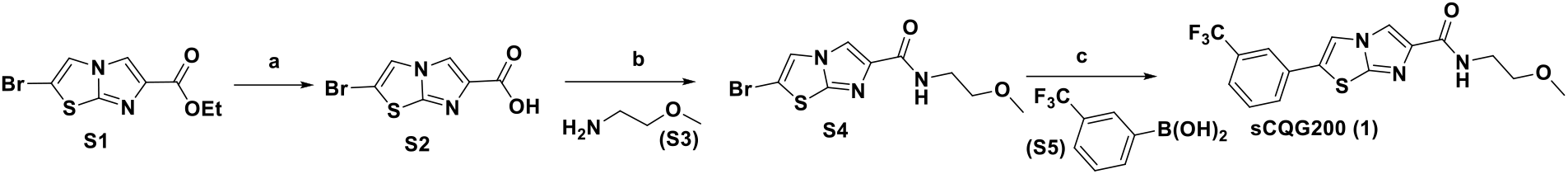
General synthetic route for **sCQG200** (**1**) and analogs. **Reagents and conditions** a) LiOH, MeOH/H_2_O, rt, 2h; b) HATU, 2-methoxyethan-1-amine (**S3**), DIEA, DMF, rt, 8h; c) (3-(trifluoromethyl)phenyl)boronic acid (**S5**), XantPhos Pd G3, CsCO_3_, Dioxane/H_2_O, 100 ^o^C, overnight

We first examined the SAR of the aryl region. The bromo intermediate **2**, which lacks the aryl substituent, was inactive against *M. tuberculosis*, indicating that the aryl group is required for anti-TB activity. Shifting the trifluoromethyl (CF_3_) substituent on the aryl ring from the *meta* to the *para* position, as in compound **3**, resulted in a >5-fold loss of potency. We next examined the effect of introducing an additional fluorine substituent in the context of the *meta*-CF_3_ analog. Fluorination at the *meta* (**5**) or *ortho* (**6**) position relative to CF_3_ improved anti-TB activity, whereas fluorination at the *para* position resulted in complete loss of activity, suggesting that substitution at this position is not tolerated. This trend was further supported by the mono-fluoro analogs **7** and **8**. The *ortho*-fluoro analog **7** was largely inactive (MIC_90_ = 24.8 µM), whereas the *meta*-fluoro analog **8** retained activity (MIC_90_= 9.9 µM), albeit with a modest reduction in potency relative to **sCQG200**. Collectively, these results indicate a strong positional dependence of aryl substitution within this series. We next examined whether difluoro substitution was tolerated. The 2,3-difluoro analog **9** retained activity comparable to that of **sCQG200**, whereas the 2,4-difluoro analog **10** showed improved anti-TB potency, with an MIC_90_ of 1.87 µM. Compared with difluoro analog **9**, the difluoro-chloro analog **11** showed an approximately 2-fold reduction in anti-Mtb activity, suggesting that potency in this series is sensitive to both the electronic/steric properties and the positional arrangement of substituents on the aryl ring. Replacement of the phenyl ring with five-membered heteroaryl groups (**12** and **13**) resulted in complete loss of anti-Mtb activity. Most analogs exhibited little or no cytotoxicity, except for analog **6**, which showed a CC_50_ of 6.89 µM in HEK293T cells.

Encouraged by the activity of the 2,3-difluorophenyl analog **10**, we next investigated the SAR of the carboxamide region in the context of this aryl substitution pattern. We first evaluated the corresponding carboxylic acid intermediate **14**, which was inactive. Replacement of the terminal methyl ether with a hydroxyl group resulted in a substantial loss of anti-Mtb activity, with analog **15** exhibiting an MIC_90_ of 19.9 µM. Replacement of the methyl ether with a methylamino group abolished activity (**16**), while replacement of the sulfone moiety with an ether was also poorly tolerated, as exemplified by analog **17**. Extension of the methyl ether in analog **10** to an ethyl ether (**18**) improved anti-Mtb potency by approximately 2-fold, prompting further exploration of this position. The isopropyl ether analog **19** showed activity comparable to that of the methyl ether analog **10**, whereas the cyclopropyl ether analog **20** exhibited potency similar to that of the ethyl ether **18**. Further increasing the cycloalkyl ring size to cyclobutyl (**21**) and cyclopentyl (**22**) substituents resulted in additional improvements in anti-Mtb activity, with MIC_90_ values of 0.94 and 0.93 µM, respectively. Collectively, these results indicate that the terminal ether region favors hydrophobic substitution and can accommodate increased steric bulk. In contrast, increased polarity at this position was poorly tolerated, suggesting that both hydrophobicity and substituent shape are important determinants of activity in this region. Encouraged by this trend, we next explored replacement of the terminal ether motif with conformationally constrained hydrophobic groups. The representative difluorocyclobutyl analog **24** retained activity comparable to that of analog **10**, indicating that the terminal ether is not essential for activity and that alternative hydrophobic amide-tail substituents are tolerated. As observed by the aryl-modified analogs, the amide-modified analogs also showed little or no cytotoxicity.

Next, we evaluated the antimycobacterial activity of selected analogs against the *Mycobacterium tuberculosis* Erdman strain and intracellular Mtb. As shown in **Table 4**, analogs **10, 21**, and **24** exhibited improved MIC_90_ values against the Erdman strain while retaining activity in the intramacrophage assay. These compounds were further evaluated in ADME profiling studies. All three analogs exhibited good stability in mouse liver microsomes (MLM), with intrinsic clearance (CL_int_) values of 7.21, 19.89, and 11.25 µL/min/mg for analogs **10, 21**, and **24**, respectively. Analogs **21** and **24** showed high plasma protein binding (PPB) in mice, with very low unbound fractions, whereas analog **10** exhibited a higher unbound fraction of 7.6%. The higher unbound fraction of **10** may be consistent with its lower lipophilicity, as reflected by its lower cLogP (3.55) compared with analogs **21** and **24** (4.32 and 5.52, respectively). Analog **10** also showed good *in vitro* intestinal permeability, with a Caco-2 apparent permeability coefficient (P_app_, A-B) of 8.7 × 10^-6^ cm/s, whereas analog **24** exhibited substantially lower permeability (P_app_, A-B = 0.77 × 10^-6^ cm/s). A cytochrome P450 inhibition panel showed that analog **24** had no noticeable inhibitory activity against the major CYP isoforms tested (IC_50_ > 50 µM for CYP3A4, CYP2D6, CYP2C19, and CYP2C9), suggesting a low potential for CYP-mediated drug–drug interactions. Analog **24** showed moderate inhibition of the human Ether-a-go-go-related Gene (hERG) potassium channel, with an IC_50_ of 8.26 µM; this liability will be monitored and addressed during further SAR optimization.

**Table 4.**
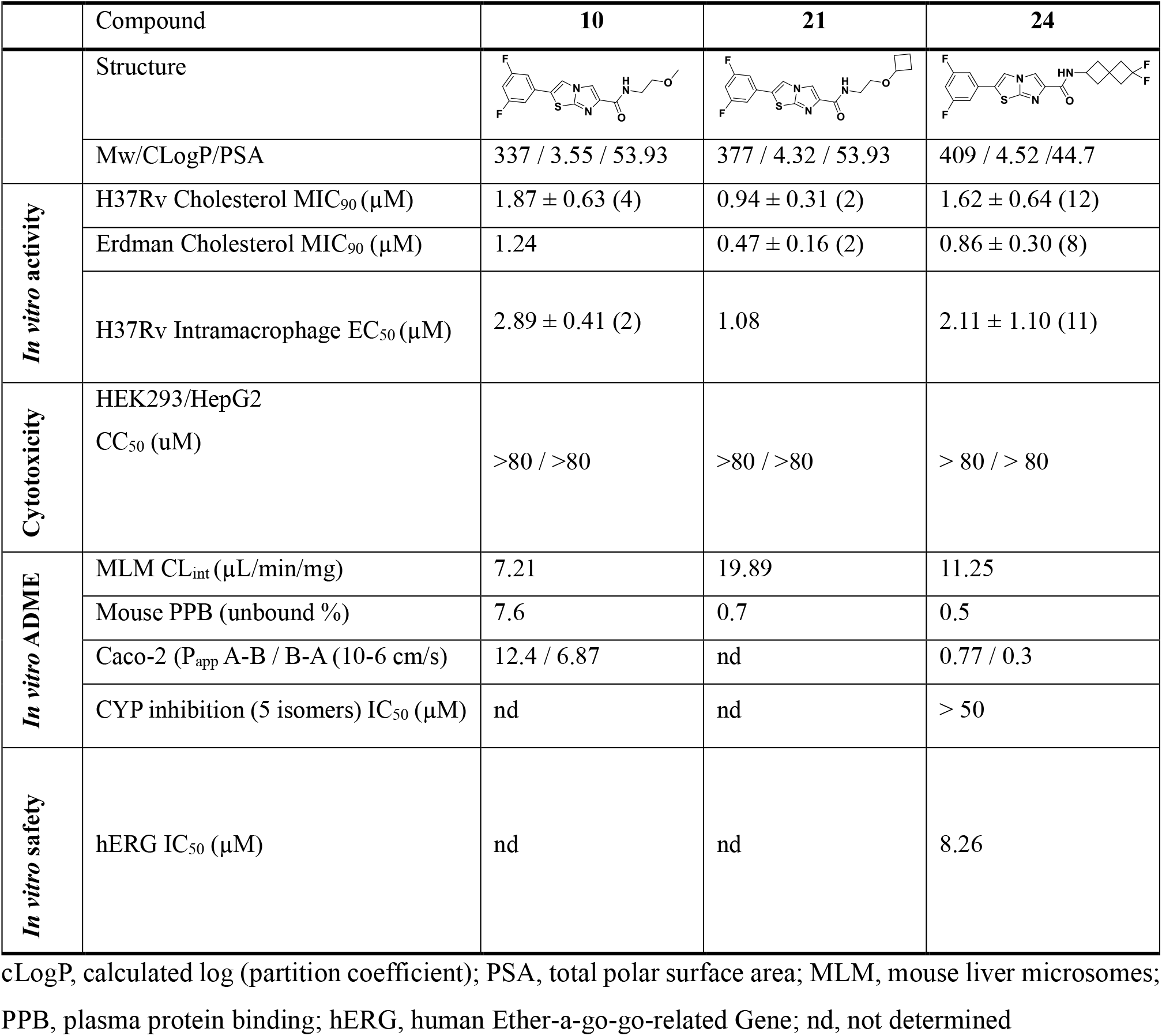
Profiling of lead compounds **10, 21** and **24**.

|  | Compound | **10** | **21** | **24** |
| --- | --- | --- | --- | --- |
|  | Structure |  |  |  |
|  | Mw/CLogP/PSA | 337 / 3.55 / 53.93 | 377 / 4.32 / 53.93 | 409 / 4.52 / 44.7 |
| *In vitro* activity | H37Rv Cholesterol MIC<sub>90</sub> (μM) | 1.87 ± 0.63 (4) | 0.94 ± 0.31 (2) | 1.62 ± 0.64 (12) |
|  | Erdman Cholesterol MIC<sub>90</sub> (μM) | 1.24 | 0.47 ± 0.16 (2) | 0.86 ± 0.30 (8) |
|  | H37Rv Intramacrophage EC<sub>50</sub> (μM) | 2.89 ± 0.41 (2) | 1.08 | 2.11 ± 1.10 (11) |
| Cytotoxicity | HEK293/HepG2<br>CC<sub>50</sub> (uM) | >80 / >80 | >80 / >80 | > 80 / > 80 |
| *In vitro* ADME | MLM CL<sub>int</sub> (μL/min/mg) | 7.21 | 19.89 | 11.25 |
|  | Mouse PPB (unbound %) | 7.6 | 0.7 | 0.5 |
|  | Caco-2 (P<sub>app</sub> A-B / B-A (10<sup>-6</sup> cm/s) | 12.4 / 6.87 | nd | 0.77 / 0.3 |
|  | CYP inhibition (5 isomers) IC<sub>50</sub> (μM) | nd | nd | > 50 |
| *In vitro* safety | hERG IC<sub>50</sub> (μM) | nd | nd | 8.26 |

cLogP, calculated log (partition coefficient); PSA, total polar surface area; MLM, mouse liver microsomes; PPB, plasma protein binding; hERG, human Ether-a-go-go-related Gene; nd, not determined

To further characterize the pharmacokinetic properties of selected analogs, compounds **10, 21**, and **24** were evaluated in mice following intravenous (IV, 3 mg/kg) and oral (PO, 20 mg/kg) administration with a solution formation with 75% PEG and 25% D5W (**Table 5**). Following intravenous administration, compound **10** was cleared rapidly (CL = 50.6 mL/min/kg) with a short terminal half-life (T_1/2_ = 0.62 h). Compound **21** showed a >2-fold reduction in clearance (CL = 20.1 mL/min/kg) and a correspondingly longer half-life (T_1/2_ = 1.42 h). Compound **24** exhibited the most favorable profile, with clearance reduced ∼15-fold relative to **10** (CL = 3.48 mL/min/kg) and a markedly prolonged half-life (T_1/2_ = 19.1 h). The steady-stage volume of distribution (Vd_ss_) values were 2.50, 1.42, and 5.55 L/kg for **10, 21**, and **24**, respectively, indicating distribution beyond the vascular compartment. Following oral administration, compound **21** demonstrated substantially improved oral exposure compared with **10**, with a C_max_ of 1791 ng/mL, AUC_last_ of 9807 h·ng/mL, and oral bioavailability of 62.1%, while maintaining a MRT_last_ of 4.74 h. Compound **24** showed the highest systemic exposure (AUC_last_ = 16,701 h·ng/mL) and a prolonged MRT_last_ of 12.8 h, although its oral bioavailability was lower (F = 29.5%) and absorption was delayed (T_max_ = 8 h). Overall, these data demonstrate substantial improvements in systemic exposure and duration of exposure for compounds **21** and **24**, providing a favorable PK basis for further evaluation.

**Table 5.**
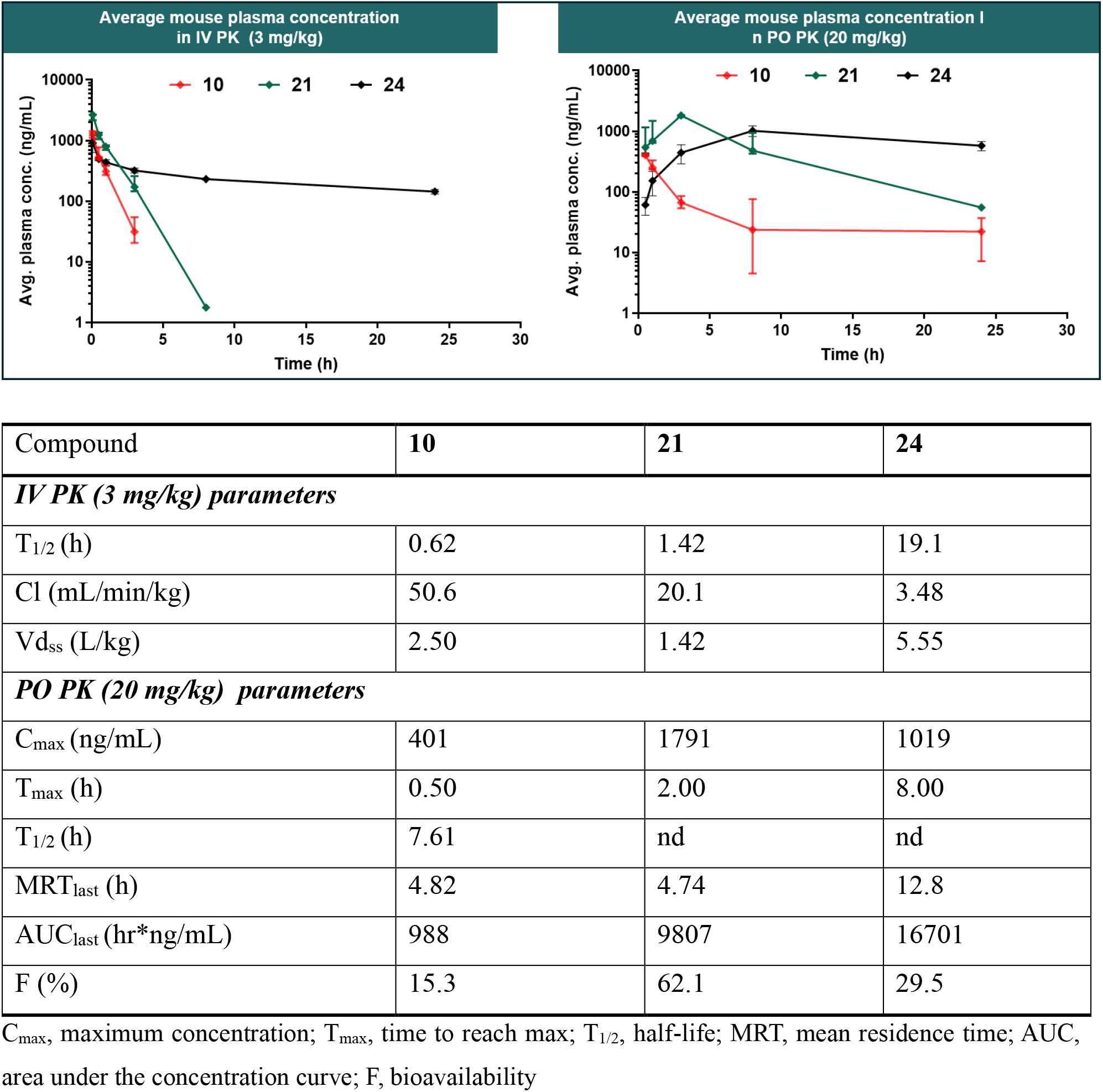
Pharmacokinetic properties of **10, 21** and **24** in mice with IV/PO administrations.

In conclusion, screening of our in-house CRESTdb small-molecule library identified sCQG200 as an imidazothiazole carboxamide with activity against *Mycobacterium tuberculosis*, including intracellular and drug-resistant strains. Retention of activity against Q203-resistant QcrB mutants suggests a potentially distinct resistance profile from established QcrB inhibitors. SAR optimization led to improved antimycobacterial potency and enabled characterization of key developability properties, including cytotoxicity, hERG activity, ADME, and pharmacokinetic properties. This structurally differentiated chemotype provides a promising starting point for further optimization to improve potency and pharmacokinetic properties while mitigating developability liabilities, including hERG activity observed for compound **24**. Studies to elucidate the mechanism of action of this series are also ongoing and will be reported in due course.

## Notes

The authors declare no competing financial interest.

## Acknowledgements

We are grateful to the Calibr compound management and pharmacology group for their assistance with this project. Special thanks to Sharon Irelan for her assistance in project management. This work was supported by a grant from the Bill & Melinda Gates Foundation #OPP1208899 to Calibr.

